# HIF-stabilization prevents delayed fracture healing

**DOI:** 10.1101/2020.07.02.182832

**Authors:** Annemarie Lang, Sarah Helfmeier, Jonathan Stefanowski, Aditi Kuppe, Vikram Sunkara, Moritz Pfeiffenberger, Angelique Wolter, Alexandra Damerau, Shabnam Hemmati-Sadeghi, Jochen Ringe, Rainer Haag, Anja E. Hauser, Max Löhning, Carsten Perka, Georg N. Duda, Paula Hoff, Katharina Schmidt-Bleek, Timo Gaber, Frank Buttgereit

## Abstract

The initial phase of fracture healing decides on success of bone regeneration and is characterized by an inflammatory milieu and low oxygen tension (hypoxia). Negative interference with or prolongation of this fine-tuned initiation phase will ultimately lead to a delayed or incomplete healing such as non-unions which then requires an effective and gentle therapeutic intervention. Common reasons include a dysregulated immune response, immunosuppression or a failure in cellular adaptation to the inflammatory hypoxic milieu of the fracture gap and a reduction in vascularizing capacity by environmental noxious agents (e.g. rheumatoid arthritis, smoking). The hypoxia-inducible factor (HIF)-1α is responsible for the cellular adaptation to hypoxia, activating angiogenesis and supporting cell attraction and migration to the fracture gap. Here, we hypothesized that stabilizing HIF-1α could be a cost-effective and low-risk prevention strategy of fracture healing disorders. Therefore, we combined a well-known HIF-stabilizer – deferoxamine (DFO) – and a less known HIF-enhancer – macrophage migration inhibitory factor (MIF) – to synergistically induce improved fracture healing. Stabilization of HIF-1α enhanced calcification and osteogenic differentiation of MSCs *in vitro*. *In vivo*, the application of DFO with or without MIF during the initial healing phase accelerated callus mineralization and vessel formation in a clinically relevant mouse-osteotomy-model in a compromised healing setting. Our findings provide support for a promising preventive strategy towards bone healing disorders in patients with a higher risk due to e.g. delayed neovascularization by accelerating fracture healing using DFO and MIF to stabilize HIF-1α.

## Introduction

Fracture healing combines temporal and spatial fine-tuned and tightly regulated regenerative processes, which lead to a complete restoration of the broken bone without scar formation. However, a minimum of 10 % of patients with fractures suffer from fracture healing disorders such as delayed or incomplete healing (non-unions) leading to immobility, pain, a loss in quality of life, and generate an economic burden for the society (*1, 2*). While trauma severity and location can determine the healing success (*3*), several risk factors have been described to potentially impair the fracture healing process such as age and lifestyle including obesity, alcohol abuse, and smoking (*4*). Of note, smoking is supposed to reduce vessel formation and to stimulate adverse immune reactions (*5*). Furthermore, chronic inflammatory diseases, such as rheumatoid arthritis (RA) or systemic lupus erythematosus, have been related to fracture healing disorders (*6*–*8*). Patients with fracture healing disorders often require several further revision surgeries. Apart from surgical intervention, local delivery of recombinant human (rh)BMP-2 into the fracture gap has been demonstrated in clinical studies to be effective (*9, 10*). However, several adverse effects in humans strongly restrict the clinical implementation of this approach (*11*) and alternative strategies or preventive measures are lacking.

If a fracture is stabilized such that inter-fragmentary movements can occur, secondary bone healing is initiated leading to bone regeneration via an endochondral ossification process bridging the fracture gap. Endochondral bone healing can be divided into five phases: i) initial pro-inflammatory phase, ii) anti-inflammatory phase; iii) fibrocartilaginous or soft callus phase; iv) woven bone or hard callus phase; and v) the remodeling phase leading to the bone restitution in form and function according to the mechanical strain (*12*). The shift from pro- to anti-inflammatory phase is a requirement for angiogenic and pro-osteogenic processes during the initial phase of fracture healing and determine the subsequent regeneration cascades (*6, 13*). This shift is essential for the initiation of bone reconstruction involving the recruitment of i) monocytes/macrophages, clearing the inflammatory scene and paving the path for revascularization, ii) pre-osteoblasts and mesenchymal stromal cells (MSCs) as basic component of bone reconstruction, iii) fibroblasts, which are required for early callus formation and bone matrix formation, and iv) endothelial cells (ECs) for neovascularization (*14, 15*). Crucial elements for successful bone healing are balanced control and termination of the pro-inflammatory cascade (*16*), proper mesenchymal differentiation and cartilage formation, controlled invasion of vessels (*17*) as well as a sufficient mechanical stabilization (*18*–*21*).

The hypoxic and inflammatory conditions in the fracture hematoma result from the disruption of vessels, accumulation of inflammatory cells, increased cell death of e.g. erythrocytes and the lack of nutrients, oxygen, high lactate and low pH – a cytotoxic environment which has to be down-regulated to maintain regenerative cells. The oxygen tension within the fracture site is reduced over the first week after trauma being accompanied by a reduction (50%) of blood flow (*22*–*27*). Therefore, cellular adaptation mechanisms towards hypoxia are strongly activated – such as the hypoxia inducible factor (HIF) signaling pathway. Both, HIF-1 and HIF-2 are essential for cells to survive hypoxic conditions and to aim at increasing the oxygen supply while reducing oxygen consumption (*28*). While HIF-1β is constitutively expressed, HIF-1α is oxygen-dependently activated and stabilized at less than 5% oxygen (*29*). HIF-1α is then translocated to the nucleus, where it heterodimerizes with HIF-1β, binds to its target sequences (hypoxia-responsive elements), and activates genes necessary for cellular hypoxic adaptation (*29, 30*). Under normoxic conditions, HIF-1α is hydroxylized by the oxygen- and iron-dependent prolyl-hydroxylase domain (PHD) enzyme/protein and degraded by cellular proteasomes.

We have previously examined fracture hematomas obtained from immunologically restricted patients and found reduced osteogenic differentiation due to reduced *runt-related transcription factor 2 (RUNX2)* expression, exaggerated immune reactions (*interleukin IL-8, C-X-C chemokine receptor type 4 CXCR4*), and high expression of *HIF1A* but inadequate expression of target genes (*31*). We also found higher numbers of monocytes/macrophages, natural killer T (NKT) cells and activated T cells within fracture hematomas of immunologically restricted patients accompanied by higher levels of IL-6, IL-8, tumor necrosis factor (TNF)α and chemokines (e.g. Eotaxin) (*32*). In order to increase target gene expression, HIF-1α can be chemically stabilized by different factors, which either inhibit the O_2_-sensing PHD such as deferoxamine (DFO) or directly interfere with the downstream effects after translocation to the nucleus e.g. the macrophage-migration inhibitory factor (MIF) (*33, 34*).

Here, we initially performed a single center retrospective study to investigate the risk factors for fracture healing disorders in a Charité-located patient cohort and to determine the clinical need to preventively support and accelerate fracture healing. Furthermore, we comprehensively systematically reviewed the available literature to delineate the potential of our drafted therapeutic approach of promoting fracture healing using DFO. To this end, we summarized several studies which demonstrated the efficacy of DFO to promote bone fracture healing in a variety of animal models (mouse, rat, rabbit) focusing on different kinds of bone defects. Experimentally, we conducted *in vitro* studies on osteogenesis, which supported the enhancing effect of HIF-1α stabilization on osteogenic differentiation of MSCs and its counteracting efficacy against e.g. glucocorticoid (GC)-induced inhibition of osteogenesis. Moreover, we tested the combination of MIF and DFO in a mouse-osteotomy-model of compromised bone healing conditions in order to evaluate their preventive capability to counteract delayed bone healing in this clinically relevant model. Thus, our study provides evidence for a promising preventive strategy to accelerate fracture healing by applying potent HIF-stabilizers during initial fracture treatment in patients at risk that may finally help to minimize bone healing disorders.

## Results

### Fracture healing disorders in a single-center patient cohort – A retrospective study

Fracture healing disorders are associated with the incidence of several different risk factors e.g. age, gender, fracture location, comorbidities of medications. Therefore, we investigated the main risk factors for fracture healing disorders in a single-center retrospective study at the Center for Musculoskeletal Surgery, Charité-Universitätsmedizin Berlin to get a more precise view of the patient’s need for a therapeutic support and acceleration of fracture healing. To this end, we screened data from inpatients treated in the hospital during 2012 (**fig. S1**). Finally, 79 cases fulfilling inclusion criteria were included in the study (**table S1**), as well as 178 controls matched for age and fracture location while patients aged < 18 years, having open fractures or metastases close to the fracture location were excluded.

First, we performed a descriptive statistical analysis to compare the two groups based on the collected parameters such as body mass index (BMI), gender, alcohol abuse (repeated consumption of alcohol per week), smoking, glucocorticoid and NSAID treatment as well as diagnosed comorbidities as RA, osteoporosis, arterial hypertension and diabetes type 2 (**table S2**). The average BMI of the two groups was similar. Comparing the control and case group, we found more male patients (53.2% vs. 44.9%) than female patients (46.8% vs. 55.1%) to be affected by fracture healing disorders. Interestingly, alcohol abuse was more often present in the control group as compared to the case group (14.4% vs 2.7%) which was the opposite for smoking (28.1% vs. 37.8%). Regarding medications, a higher number of patients with fracture healing disorders were continuously treated with glucocorticoids (6.3% vs. 1.6%) and NSAIDs (7.6% vs. 2.7%) as compared to controls. Although the frequencies of osteoporosis and diabetes type 2 were comparable within both groups, the incidences of RA (6.3% vs. 0.5 %) and arterial hypertension (43% vs. 33.5%) were higher in the case group. To select potentially relevant factors, an univariable logistic regression was performed (**Table 1**) using a significance level of *0.15* resulting in the selection of smoking, RA and arterial hypertension for detailed analysis via multivariable logistic regression. Age and gender, parameters well-known to be associated with a poor fracture healing outcome, were additionally included in the subsequent multivariable logistic regression. Statistical analysis using multivariable logistic regression showed a high significance for RA (*P* = 0.028) and a trend for smoking (*P* = 0.075) to be associated with fracture healing disorders such as non-unions.

**Table 1:** Univariable and multivariable logistic regression.

|  | *Univariable* |  | *Multivariable* |  |  |  |
| --- | --- | --- | --- | --- | --- | --- |
|  | Omnibus test | P-value | 5 | 4 | 3 | 2 |
| Age | 0.597 | 0.597* | 0.118 | 0.202 | - | - |
| Gender | 0.219 | 0.219* | 0.148 | - | - | - |
| BMI | 0.212 | - |  |  |  |  |
| Alcoholism | 0.657 | - |  |  |  |  |
| **Smoking** | **0.131** | **0.129** | **0.065** | **0.063** | **0.048** | **0.075** |
| **Rheumatoid Arthritis** | **0.006** | **0.022** | **0.057** | **0.047** | **0.040** | **0.028** |
| Glucocorticoids | 0.725 | - |  |  |  |  |
| NSAIDs | 0.336 | - |  |  |  |  |
| Osteoporosis | 0.813 | - |  |  |  |  |
| **Arterial Hypertension** | **0.143** | **0.142** | **0.085** | **0.058** | **0.137** | - |
| Diabetes Type 2 | 0.908 | - |  |  |  |  |

### Deferoxamine as potent target for fracture healing disorders – A systematic literature review

We have previously found that immunologically restricted patients (including e.g. autoimmune disease) show a disturbed response to hypoxia in the fracture hematoma (*31*). Therefore, we hypothesized that HIF-stabilization can accelerate fracture healing in those patients. The inhibition of iron-dependent PHDs by iron chelators or competitors activates HIF-mediated pathways such as angiogenesis and osteogenesis. The iron chelator DFO is well-known from *in vitro* and *in vivo* studies to stabilize HIF (*35, 36*).

Given this background, we performed a systematic literature review to delineate existing preclinical studies on the effectiveness and efficiency of DFO in fracture healing. In detail, we asked the question whether the local application of DFO in the fracture gap enhanced bone formation (μCT; histomorphometry) during fracture healing in animal models with normal or disturbed fractures of long-bones or *Ossa irregularia* (*mandibula* or *zygomatic arch*). The complete search strategy can be found in figure S2 and S3. We included 20 studies for a descriptive analysis. A meta-analysis was not applicable due to the variability of studies, the variety of models and the information provided (e.g. data values not given; confidential intervals not indicated). Nevertheless, all included studies demonstrated the efficacy of DFO to promote bone fracture healing in a variety of animal models (mouse, rat, rabbit) with different bone defects (**Table 2**). In these studies, DFO was applied in varying doses (in mouse: 20 μl of 200 μM – 400 μM) either once or repetitively by local injection directly into the bone defect/gap or by loading onto a scaffold implanted into the bone defect/gap to counteract a delayed healing or non-union. DFO treatment resulted in a strong promotion of angiogenesis/vessel formation and bone regeneration independent of the species, model and evaluation methods (22–39).

**Table 2:**
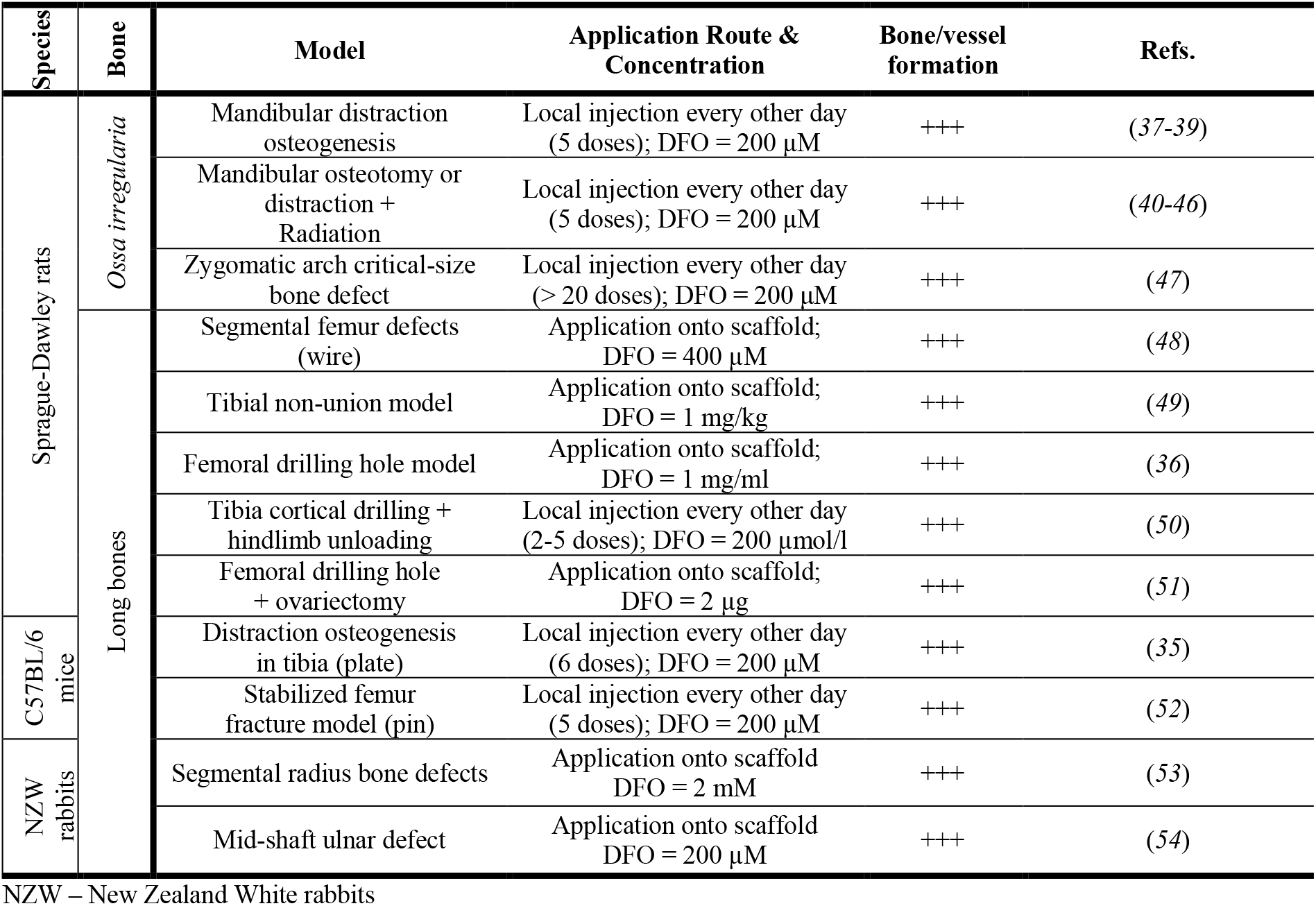
Systematic literature review on the potential of DFO to accelerate bone formation/healing.

Based on the published data, we concluded that a single injection of DFO might be sufficient to enhance fracture healing under compromised conditions. In addition, we asked if the effect of DFO can be increased by combination with MIF which we have shown to further enhance HIF activity, especially in immune cells and endothelial cells (*55*).

### Combining DFO and MIF to enhance in vitro calcification of hMSCs

To evaluate the potential of MIF and DFO as enhancer of bone formation *in vitro*, different concentrations alone and in combination were tested in an osteogenic differentiation assay of bone marrow derived human (h)MSCs. High concentrations of Dexamethasone (Dex; 10^−3^ M) were used as a technical *in vitro* model to strongly induce delayed calcification while 10^−8^ M Dex was the respective control which is usually included in the osteogenic medium (OM). Large differences in calcification were observed after 4 weeks under normoxic conditions and used further titration experiments (**fig. S4**; **fig. S5**; **Fig. 1**). Normoxic conditions represented the inadequate adaptation to hypoxia as mentioned before. Varying concentrations of MIF and DFO alone and in combination were examined for their effect on hMSC calcification. The application of DFO alone showed significant increases in calcification after 4 weeks at 62.5, 125, 250 and 500 μM which was also observed at 250 and 500 ng/ml MIF (**fig. S5A, B**). A double normalization to both controls (10^−3^ M and 10^−8^ M Dex) revealed that MIF alone did not enhance calcification (**Fig. 1A**), while DFO alone already showed mean increases between 43% - 45.9% (**Fig. 1B**). The combination of MIF and DFO led to a significant increase in calcium deposition when using 50 ng/ml MIF + 125 μM DFO and 100 ng/ml MIF + 125 μM DFO with a mean increase of 61.7% to 86.7% for both groups (**fig. S5C**; **Fig. 1C**).

**Figure 1:**
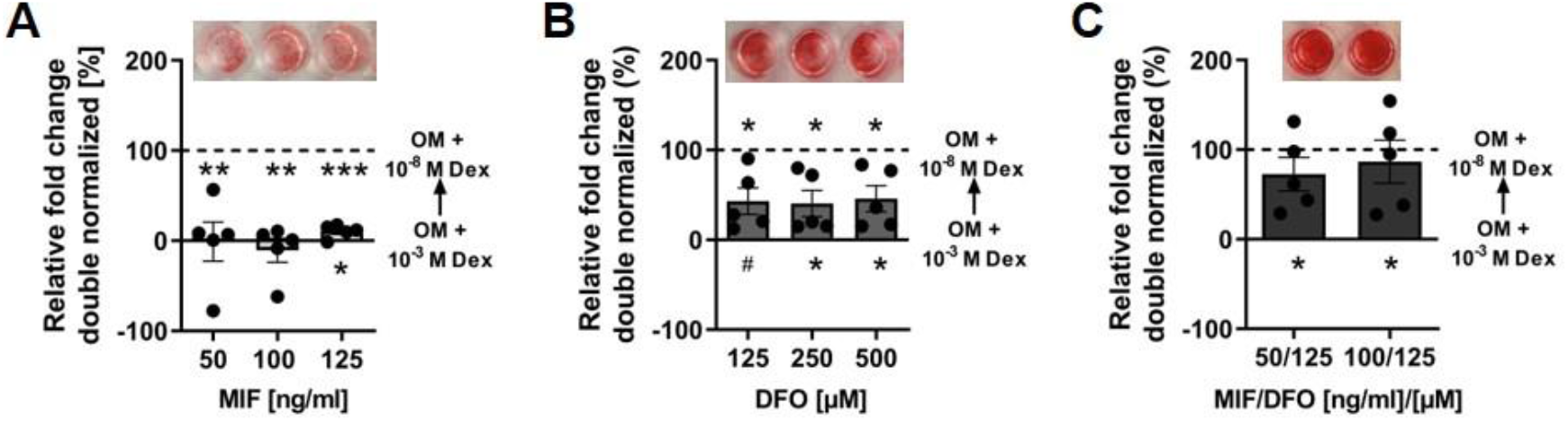
In vitro studies on the effect of MIF and DFO on hMSC calcification. Double normalization of OD values (Dex 10^−3^ M = 0 %; Dex 10^−8^ M = 100 %) for selected MIF (A), DFO (B) and MIF/DFO (C) concentrations. Asterisks above and below the bars indicate significant differences as compared to the respective control. Bar graphs show mean ± SEM and individual data points. One sample t-test was used to determine the statistical significance; P-values are indicated with ^#^P < 0.07; *P < 0.05; **P < 0.01; ***P < 0.001. Exemplary images of Alizarin red staining in 96 well are displayed before quantification.

### The effect of MIF and DFO on in vivo bone formation in a delayed healing model

Based on the results from the systematic review and our *in vitro* experiments, we tested MIF and DFO, both alone and in combination in an experimental delayed healing model using a modified mouse-osteotomy which results in a local blockage of angiogenesis and results in a delayed bone healing due to insertion of an absorbable bovine Col-I scaffold (ACS) in the osteotomy gap (*56*). We previously demonstrated that this model features a disturbance in cell invasion, vessel formation and consecutively bone formation when compared to empty-gap controls at 2 and at 3 weeks after osteotomy (*56*). The osteotomy gap (0.7 mm) was introduced in the femur of 12 weeks old female C57BL/6N mice. A stable fixation of the osteotomized bone was obtained using an external fixator.

Bone formation was significantly intensified in the DFO and MIF/DFO treated groups as measured by bone volume fraction at 3 weeks post-osteotomy (**Fig. 2A**; **fig. S6A**). We confirmed our finding by histological evaluation quantifying Movat’s pentachrome staining (**Fig. 2B, C**; **fig. S6B, C**). As a result, we observed increased levels in total mineralized bone tissue in the fracture area and in the fracture gap in all treatments groups after 2 weeks (**Fig. 2C**). Interestingly, MIF also significantly induced total bone formation especially in the fracture gap at 2 weeks when compared to the corresponding ACS control, while in the other groups mineralized bone formation was more pronounced within the gap (**Fig. 2C**). Of note, we observed a higher cartilage content in the MIF and DFO group at 2 weeks as compared to the ACS control, while MIF/DFO exhibited similar amounts of cartilage. These differences were not present at 3 weeks, since animals with only ACS show a delayed endochondral ossification (between week 2 and 3; **fig. S6C**) (*56*). However, the complete bridging of the fracture gap with cartilage and mineralized bone was observed in 50% of DFO and MIF/DFO mice additionally indicating an acceleration of the endochondral ossification process (**fig. S7**). Interestingly, bridging between cortices was more often observable in the MIF/DFO treated group (50%) (**fig. S7**). Finally, DFO and MIF/DFO led to a higher recruitment of osterix (Osx)^+^ osteoprogenitors/osteoblasts to the fracture gap at 2 weeks (**Fig. 2D, E**). In addition, Osx^+^ cells were more present at day 3 in the DFO group and day 7 in MIF group (**fig. S6D**). Taken together, these data suggest that both DFO and MIF/DFO accelerated the endochondral ossification process during the fracture healing in our delayed healing mouse-osteotomy-model.

**Figure 2:**
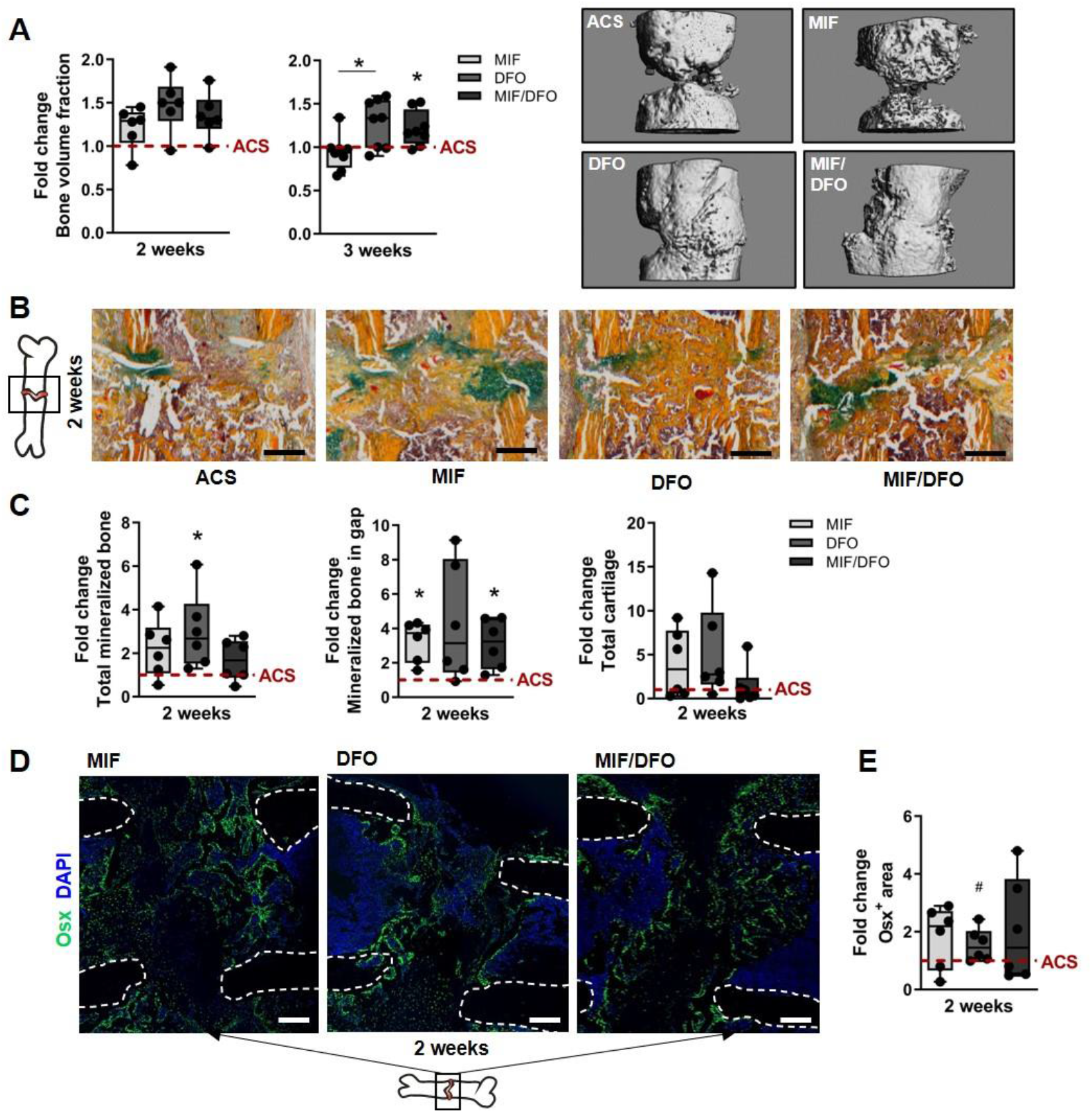
Bone regeneration in a delayed healing model after single dose of MIF or/and DFO. (A) MicroCT quantification at 2 weeks and at 3 weeks post-osteotomy normalized to the median of the ACS group (indicated as dotted line = 1). Bone volume fraction = bone volume/callus volume. Representative 3D microCT reconstructions at week 3. (B) Representative images of Movat’s pentachrome staining for each group at week 2. yellow – mineralized bone/scaffold; green – cartilage; magenta – bone marrow. (C) Histomorphometry of Movat’s pentachrome staining using ImageJ. Data were normalized to the median of the ACS group (indicated as dotted line = 1). (D) Representative images of immunofluorescence staining of Osterix (Osx) and its quantification. White dotted lines indicate cortices. Schematic bone indicates alignment of images. Scale bars indicate 200μm. Data are shown as box plots with the median as horizontal line, interquartile range as boxes, minimum/maximum as whiskers and individual data points. Wilcoxon signed rank test was applied to determine difference against the ACS control group (hypothetical value = 1) and Kruskal Wallis test with Dunn’s multiple comparison test was used to compare groups. ^#^P < 0.07; *P < 0.05.

### Vessel formation is increased by HIF-stabilization

Revascularization is crucial for bone regeneration, tightly regulated by the microenvironment and appears in two waves at day 7 and day 21 (*17, 57*). CD31^+^ endothelial progenitors enter the fracture gap during the initial phase of fracture healing (until day 7) (*57*). DFO is known to strongly promote revascularization by the induction of vascular endothelial growth factor (VEGF) expression, a target gene of HIF-1α (*34*). Therefore, we analyzed the osteotomy site in our delayed healing model for the presence of CD31^+^ Emcn^+^ vessels. At week 2 and 3, we found significantly more CD31^+^ Emcn^+^ vessels in the fracture gap of the treatment groups compared to the ACS control group while cell invasion was more pronounced at week 2 and comparable to the ACS control group at week 3 (**Fig. 3A-C**). Pixel intensity analysis revealed elevated Emcn and CD31 expressions in the DFO and MIF/DFO group at week 2 indicating a higher appearance of CD31^+^ and Emcn^+^ cells and a higher vascular formation (**Fig. 3D-F**). Interestingly, MIF alone also induced the expression of CD31 in the fracture gap at week 2 compared to the ACS group (**Fig. 3E**). However, expression levels were comparable between all groups at week 3. When examining earlier timepoints (day 3 and 7), we observed a comparable appearance of CD31^+^ endothelial progenitors in all groups but the DFO group (**fig. S8A**). In the DFO group, we observed a reduced number of CD31^+^ endothelial progenitors at day 7, which was in line with the overall low cell number (DAPI) in the fracture gap during the early stage (**fig. S8A**). Moreover, immunofluorescence images indicated a pronounced invasion of CD31^+^ endothelial progenitors in the region adjacent to the fracture gap (**fig. S8B**). We conclude from our data that MIF, DFO as well as the combined MIF/DFO enhanced and accelerated revascularization to a considerably higher extent than the corresponding control group as seen at week 2 and 3.

**Figure 3:**
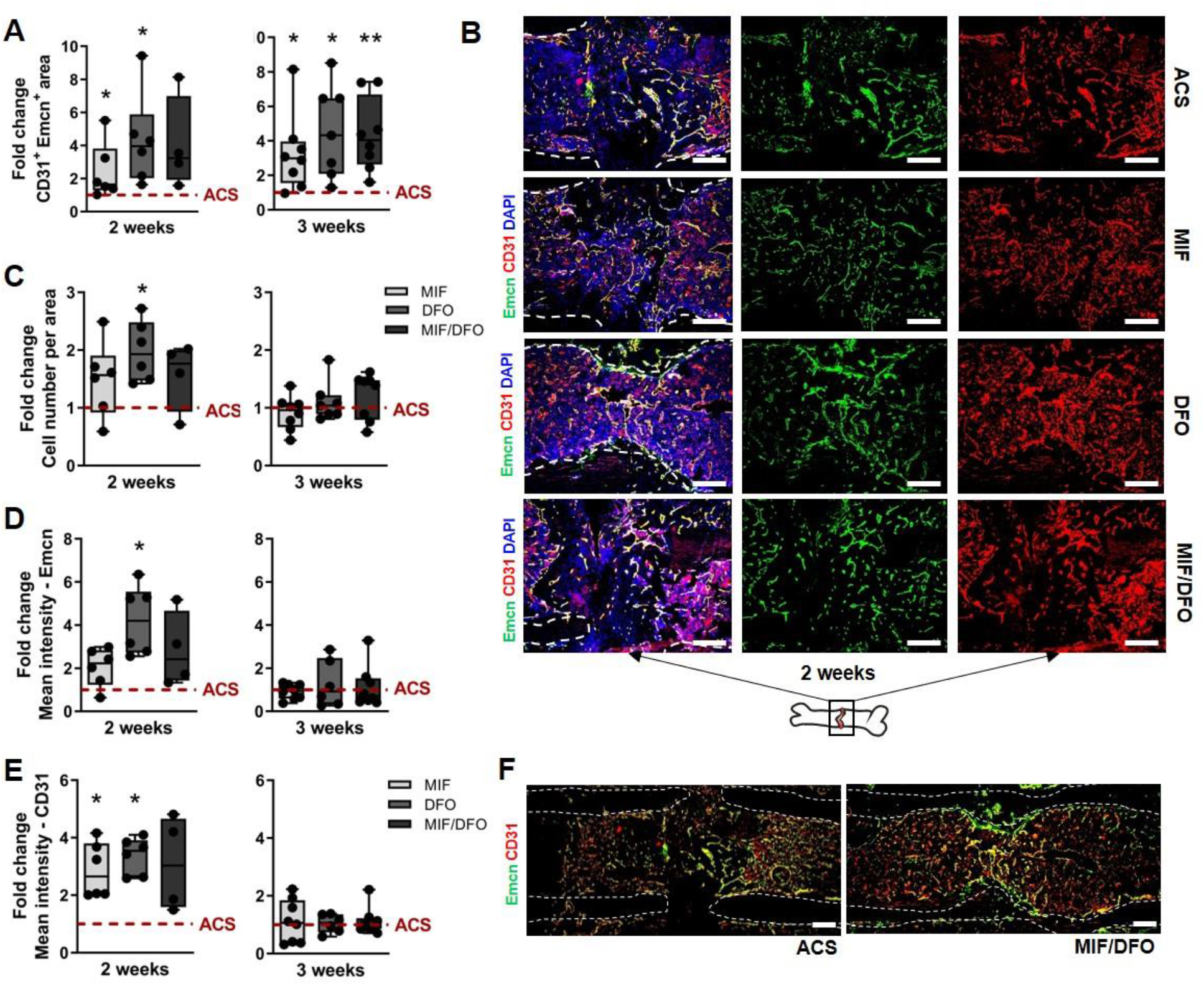
Revascularization in a delayed healing model under MIF, DFO and MIF/DFO treatment. (A-C) Quantified CD31^+^Emcn^+^ stained areas (A) and cell numbers per area (C) normalized to the median of the ACS group (indicated as dotted line = 1) and (B) corresponding representative images for week 2 and 3 (N = 6-8). (D, E) Pixel based intensity analysis of Emcn (D) and CD31 (E) in the fracture gap normalized to the median of the ACS group (indicated as dotted line = 1) and (F) representative images of the combined staining for ACS and MIF/DFO at week 2. (N = 6-8). White dotted lines indicate cortices. Schematic bone indicates alignment of images. Scale bars = 200μm. Data are shown as box plots with the median as horizontal line, interquartile range as boxes, minimum/maximum as whiskers and individual data points. Wilcoxon signed rank test was applied to determine difference against the ACS control group (hypothetical value = 1) and Kruskal Wallis test with Dunn’s multiple comparison test was used to compare groups. *P < 0.05; **P > 0.01.

### DFO and MIF/DFO lead to enhanced presence of macrophages and TRAP^+^ cells

Macrophages are essential during fracture healing and we have previously reported the crosstalk between macrophages and vessel especially during the early phase (*56, 57*). Since the treatment with both DFO and MIF/DFO resulted in an enhanced and accelerated endochondral ossification (Fig. 2) and revascularization (Fig. 3), we further asked whether DFO and MIF/DFO contribute to an increased macrophage invasion. Therefore, we analyzed the osteotomy area for the presence of F4/80^+^ cells at 3 days, 7 days, 2 weeks and 3 weeks post-surgery. We observed a higher appearance of these cells in the DFO group at day 3 indicating a faster recruitment to the fracture gap, while at day 7 the number was observably reduced as compared to day 3 and the corresponding control (ACS) (**Fig. 4A, B**). No differences were found between the other groups. Furthermore, we observed a more pronounced presence of tartrate-resistant acid phosphatase (TRAP)^+^ cells in the DFO and MIF/DFO group at week 2 than in the ACS group, which was significantly reduced in the MIF group at week 3 when compared to the ACS and DFO group (**Fig. 4C, D**). TRAP is a well-known marker of osteoclasts but can be also found on activated macrophages (*58*). In addition, quantifications of the scaffold area after 2 and 3 weeks indicated a significant reduction of the ACS in the osteotomy gap of both the DFO group (at week 2 and 3) and the MIF/DFO group (at week 2) (**fig. S9**).

**Figure 4:**
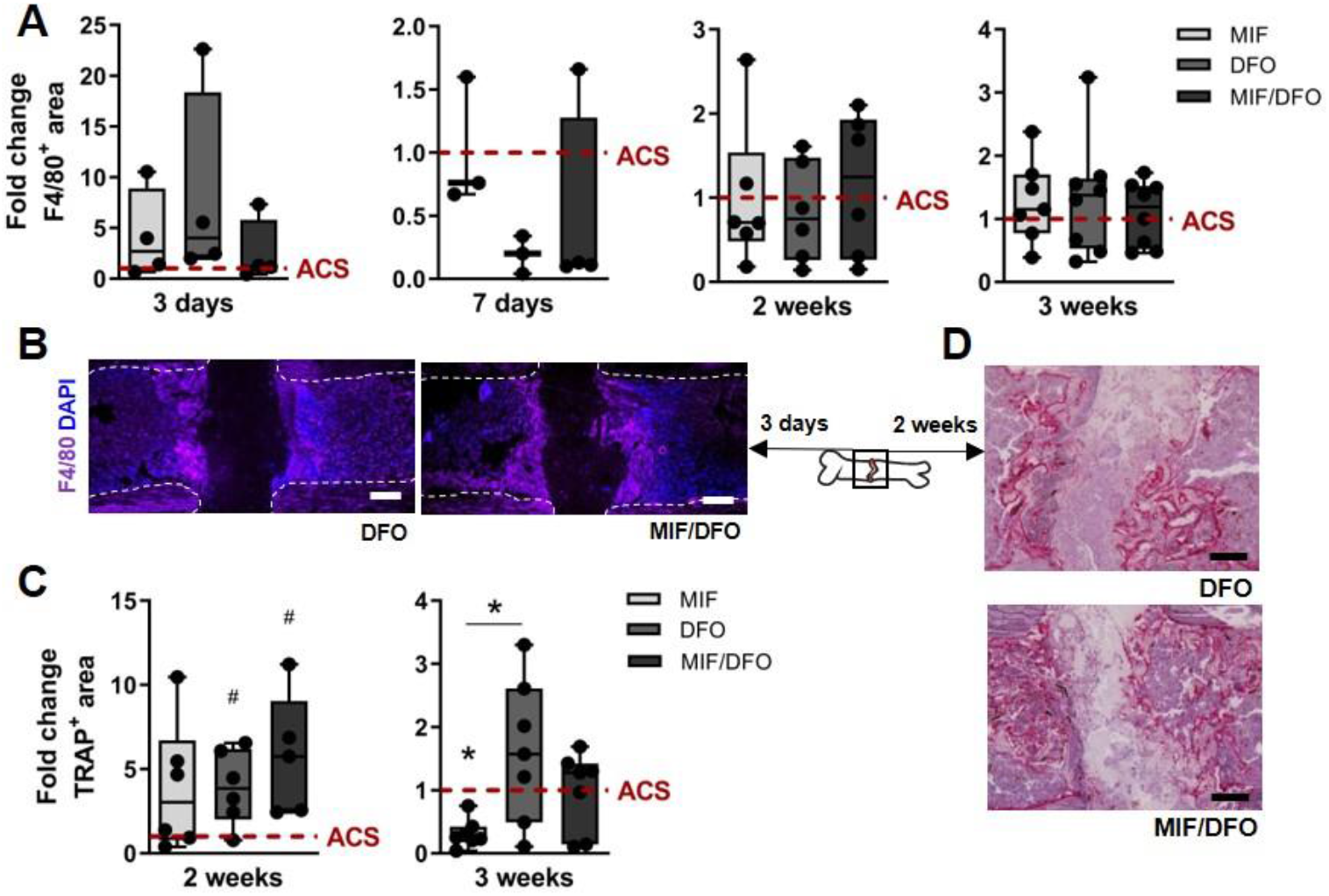
Presence of F4/80^+^ macrophages and TRAP^+^ cells within the fracture gap. (A) Quantified F4/80^+^ area in the gap after 3, 7 days and 2, 3 weeks normalized to the median of the ACS group (indicated as dotted line = 1; day 3, 7: N = 3-4; week 2, 3: N = 6-8). (B) Representative images for DFO and MIF/DFO at day 3. White dotted lines indicate cortices. Schematic bone indicates alignment of images. (C) Quantification of TRAP^+^ area at week 2 and 3 normalized to the median of the ACS group (indicated as dotted line = 1) and (D) representative images. Scale bars = 200μm. Data are shown as box plots with the median as horizontal line, interquartile range as boxes, minimum/maximum as whiskers and individual data points. Wilcoxon signed rank test was applied to determine difference against the ACS control group (hypothetical value = 1) and Kruskal Wallis test with Dunn’s multiple comparison test was used to compare groups. ^#^P < 0.07; *P < 0.05.

Together, these results suggest that both DFO and MIF/DFO promote recruitment of macrophages or local proliferation during the early phase (day 3) and a pronounced expression of TRAP (osteoclast differentiation) thereby supporting the bone regeneration process by providing the space for revascularization.

### Polyglycerol sulfate-based hydrogels as a potential releasing system for DFO

Our results provide evidence for a beneficial effect of DFO in fracture healing and in combination with MIF. In order to delineate a therapeutic option for fracture healing disorders based on HIF-stabilization for e.g. patients with a delayed healing potential due to immunological and/or angiogenic constraints (*31, 59*), we evaluated the potential of an appropriate delivery/release system. Based on the observation that the fracture hematoma obtained from immune-suppressed patients revealed an upregulated inflammatory profile during the initial phase of fracture healing, we tested dendritic polyglycerol sulfate (dPGS)-based polyethylene glycol-dicyclooctyne (PEG-DIC) hydrogels, which have been developed to act anti-inflammatory (*31, 32, 60–62*). In addition, we used non-sulfated dendritic polyglycerol (dPG)-based PEG-DIC hydrogels as control and potential alternative. Release of DFO from dPG-based hydrogels was slower compared to dPGS-based hydrogels over an observation span of 9 days (**Fig. 5A**). Supernatants from the release assay were transferred to HEK 293 cells, and protein was collected after 24h. Western blot analysis still indicated the functionality of the released DFO to stabilize HIF-1α (**Fig. 5B**). hMSCs were co-cultivated for 2 weeks with the DFO-loaded hydrogels and calcification was analyzed by Alizarin red staining (**Fig. 5C, D**). Unexpectedly, dPGS-based hydrogels seemed to have an inhibitory effect on *in vitro* hMSC calcification independent of DFO loading. Thus, we concluded that the combination of DFO with dPG-based hydrogels could be a beneficial approach regarding fracture healing. Moreover, a single dose application/injection proved to be a promising alternative.

**Figure 5:**
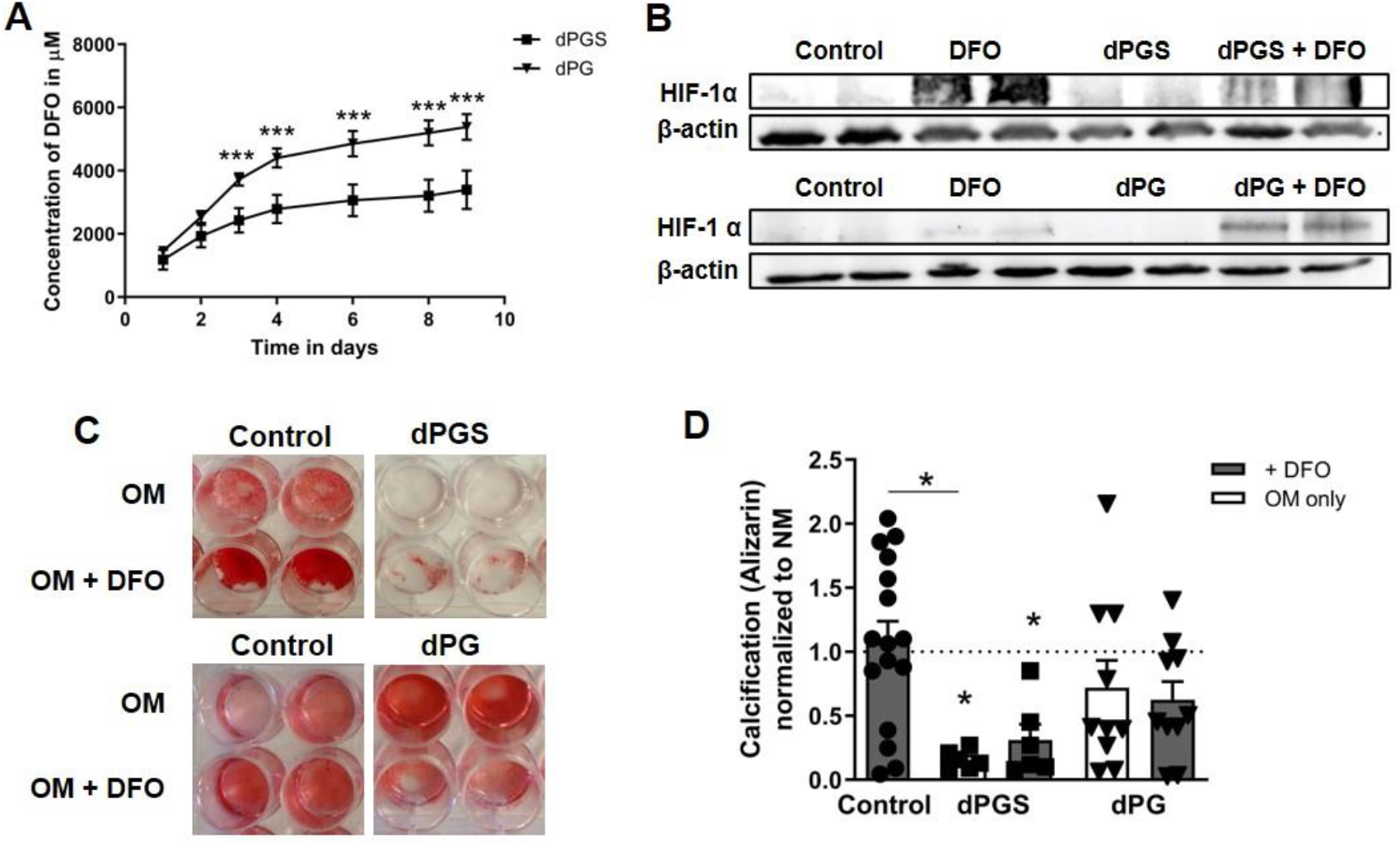
Release studies of DFO from a polyglycerol sulfate-based hydrogel (dPGS) and a polyglycerol-based hydrogel (dPG). (A) DFO release studies from the hydrogels measured over 9 days. Data is shown as mean ± SD (N = 6). Two-way ANOVA with Bonferroni posttest was performed to determine significant differences. ***P < 0.001. (B) Western blot for HIF-1α and β-actin from HEK 293 cells treated with supernatants from the release experiments. (C, D) Alizarin staining was performed to determine the effect of the hydrogels and the released DFO on hMSC calcification at 2 weeks. (N = 6-11). OM = osteogenic medium/control. Exemplary images of Alizarin red staining in 24 well are displayed. Data are shown as box plots with the mean ± SEM and individual data points. Wilcoxon signed rank test was applied to determine difference against the OM control group (hypothetical value = 1) and Kruskal Wallis test with Dunn’s multiple comparison test was used to compare groups. *P < 0.05.

## Discussion

Patients with fracture healing disorders often require further surgeries, experience substantial pain and suffer from prolonged functional impairments. While fracture healing is usually completed within 3 to 4 months, non-unions are identified if healing does not succeed after 9 months (no radiological bridging of fragments is visible; (*1, 2*). Current treatment modalities of non-union include surgical revision, autologous bone grafting or local stimulation of healing by e.g. rhBMP-2 for local delivery into the fracture gap. However, several adverse effects have been reported with BMP-2 and thus clinical usage is strongly restricted (*63*). Preventive measures for patients potentially at risk do not yet exist. Therefore, strategies that would allow to treat patients with a probable lack in their healing capability early on would be highly desirable. Our single-center retrospective study identified specifically patients with dysregulation of the immune system (e.g. RA) or disturbed capacities in angiogenesis (e.g. smoking) as being at high risk to experience a non-union (**Table 1**), which is in agreement with other reports (*6, 64*). Induction of regenerative processes within the initial phase of fracture healing depends on an adequate cellular adaptation to the hypoxic microenvironment of the fracture gap (*31*). Therefore, we hypothesized that stabilization of HIF-1α could be an effective approach for the prevention of fracture healing disorders. Based on a systematic literature review, we identified DFO as stimulatory factor in bone regeneration for both a variety of clinical bone healing settings (including mandibular defects and critical size defects) and across various species and models (mouse, rat, rabbit) (**Table 2**). *In vitro*, we could demonstrate the effect of a single-dose application of DFO alone and DFO in synergy with MIF to counteract a GC-induced inhibition of calcification during osteogenic differentiation of hMSCs. Finally, we could demonstrate in an *in vivo* proof-of-concept experiment that DFO alone or in synergy with MIF can prevent delayed bone healing in a locally impaired healing model that uses a bovine Col I scaffold in a mouse-osteotomy.

Since GCs negatively affect bone metabolism, it is not surprising that GCs have also been reported to negatively affect bone healing (*65*). Although GCs are essential for osteogenic differentiation at low concentrations, high GC concentrations inhibit osteogenic differentiation and proliferation while favoring adipogenesis (*66*–*71*). Therefore, we selected the potent GC dexamethasone to effectively inhibit *in vitro* calcification of hMSCs as an *in vitro* model for disturbed bone synthesis to test the ability of HIF–stabilizers to re-establish osteogenic differentiation within the present study (Fig. 1). Of note, beside its stabilizing effect on HIF-1α, DFO directly influences hMSC differentiation via beta-catenin signaling cascades (*72*). Interestingly, Cobalt(II) chloride, another HIF-stabilizer also increased osteogenic differentiation as recently reported (*73*). Nevertheless, the counteracting potential of DFO for GC-induced inhibition of osteogenic differentiation as shown in the present study, has not been reported, yet.

However, the ability of MIF, the natural counter-regulator of GC action, to support hypoxia-mediated HIF-1α stabilization has been shown independently by two groups (*33, 74*). The *in vitro* findings presented here demonstrate that the impact of MIF alone is not sufficient to overcome the high-dose GC–mediated suppression of osteogenic differentiation. However, MIF – in combination with DFO – synergistically enhanced the counteracting potential of DFO on hMSC calcification (**Fig. 1**). Taken together, DFO and its combination with MIF re-established osteogenic-induced hMSC calcification in a high-dose GC *in vitro* model for disturbed bone synthesis.

When analyzing HIF stabilizers/enhancers in an *in vivo* model of delayed healing using a mouse-osteotomy-model, we clearly demonstrated that application of DFO alone or in combination with MIF enhanced mineralized callus formation after 14 and 21 days (**Fig. 2**). This is in line with several previous reports on DFO administered in rat- or mouse-osteotomy-models of *Ossa irregularia* (mandibula or zygomatic arch) and long bones (femur or tibia) (*35–48, 50, 53, 54, 56, 75, 76*). All studies showed a strong positive effect of DFO on bone and vessel formation although the application routes and concentrations differed. Most comparable to our present work are the studies of Wan *et al*. and Yao *et al*. using the mouse-osteotomy-model with medullary pin fixation, but with day-wise repeated local injections of 200 μM DFO (*35, 77*). In the study presented here, our hypothesis was that a single dose of DFO alone or in combination with MIF is sufficient to accelerate bone formation. Indeed, we could verify our hypothesis by providing evidence for the enhanced mineralized bone formation at later time points (14 and 21 days; **Fig. 2**). Although the combination with MIF was supposed to further enhance the DFO-mediated enhancement of callus mineralization and bone formation, we assume that the DFO effect alone is strong enough and, therefore, masks the potential additional effect of MIF. However, comparing the histomorphometric results on mineralized bone formation endosteal or intracortical, MIF alone and in combination with DFO showed significantly more mineralized bone in the endosteal compartment after 14 days (Fig. 2) indicating a pivotal role for MIF alone during fracture healing. Ondara *et al*. described higher expression levels of MIF during the fracture healing process, which has been also described in other regenerative processes such as wound healing (*78, 79*). MIF deficient mice showed impaired fracture healing caused by a reduced number of osteoclasts and increased osteoid production (*80*). In our hands, MIF, DFO and MIF/DFO strongly enhanced revascularization much faster than in the control group as shown by the ingrowth of CD31^hi^ Emcn^hi^ ECs into the osteotomy gap (week 2 vs. 3; **Fig. 3**). Kusumbe *et al*. described these cells to be part of a bone tissue specific vessel subtype linking angiogenesis and bone formation via Notch and HIF-1 α signaling and located the CD31^hi^ Emcn^hi^ ECs to the bone surfaces and into the growth plate (*81, 82*). Moreover, stabilizing HIF by hypoxia or DFO leads to an induction of *VEGF* expression in different cell types being the major driver of vascularization also during fracture healing (*37, 52, 76*).

In addition, we observed an accelerated recruitment of macrophages and osteoclast activity based on TRAP activity in the DFO and MIF/DFO group (**Fig. 4**). DFO strongly supported resorption of biomaterial by enhancing osteoclast activity (*75*), while MIF is well-known to promote osteoclastogenesis by interacting with the RANKL pathway (*80, 83–85*). Furthermore, macrophages play a pivotal role during the whole fracture healing process. Most importantly they promote vascularization and angiogenesis by degrading ECM, which supports the release of angiogenic factors (*86, 87*). Very recently, we have demonstrated the close interconnection between macrophages and vessel formation during fracture healing (*57*).

Finally, we propose dPG as a release system to apply DFO, which may not interfere with the healing process. The advantage of synthetic, biodegradable hydrogels such as dPG is the possibility to adjust the properties of the hydrogel to the specific requirements of the fracture gap. We found that the combination of DFO with dPG could be a promising approach (Fig. 5). However, further studies are needed to optimize the delivery system by further modifications (*88*). Until today, no appropriate delivery system has been approved clinically.

In summary, our data provides convinving evidence on the potential of DFO to accelerate bone healing by enhancing mineralization and vessel formation. In addition, MIF alone used at a concentration of 100 ng/ml rather showed inhibitory properties in the regeneration process. The additional effect of MIF on top of the DFO effect was only seen for a few outcomes – e.g. vessel formation. Therefore, it can be supposed that MIF acts concentration-dependent, and further studies on the dosage finding are needed. Here, we showed that the combination of HIF-stabilizers can counteract delayed fracture healing. DFO is approved by the FDA, commercially available (e.g. Desferal® by Novartis AG) and listed on World Health Organization’s List of Essential Medicines. We consider DFO as suitable for rapid clinical translation to improve fracture healing and to be used as preventive strategy to avoid bone healing disorders in patients at high risk (e.g. RA and smoking). Therefore, we are currently striving to start a multi-centric confirmatory study with the long-term goal of clinical translation.

### Limitations

In the present study, hMSCs were isolated via migration from the bone marrow although normal protocols recommend density gradient centrifugation. We see increased cell numbers that can be an indication for higher heterogeneity in the following cell culture, which can influence the experimental outcomes. Density gradient centrifugation also has disadvantages such as the loss of smaller cell populations including high proliferative hMSCs. Moreover, there are varying protocols for density gradient centrifugation provided in the literature which does not guarantee reproducibility (*89*). Furthermore, there is strong evidence in the literature that freshly isolated hMSCs differ from isolated and cultivated hMSCs in their transcriptome and secretome indicating that conclusion from *in vitro* studies should be translated carefully towards *in vivo* assumptions (*90*). In the present study, the *in vitro* studies were rather used as a tool to get insights for further *in vivo* studies than investigating specific pathways.

For the *in vitro* studies on DFO/MIF in our GC-induced delayed calcification assay, all hMSCs were expanded and cultivated in monolayer under normoxic condition which does not parallel the normal bone marrow niche, particularly 3D and hypoxia (*91, 92*). Moreover, a heterogenic population of hMSCs was used for the assays while distinct subpopulations can be influenced differently by the treatments (*93*). In addition, high dexamethasone concentrations were required to mimic significant inhibitory effects of GCs *in vitro*. Those concentrations do not resemble clinically used dosages.

Additionally, it should be taken into account that the present study was conducted in mice, and the interpolation to the human is limited. In general, in orthopedic research rodents as well as large animal models are most commonly used. Mice are favored for basic research questions due to the possibility of genetical modifications. In contrast, sheep or pigs are preferred for translational approaches, and rats are more often used for pharmacological interventions and toxicological studies. Most animal species show slight analogies to the human bone macro- and microstructure. Main differences between mice and humans comprise permanent opening of the growth plate in the epiphyses of long bones leading to a lifelong skeletal modeling, the lack of a Haversian system and low cancellous bone content at the epiphyses of long bones (*94, 95*). Here, a mouse-osteotomy-model was used which does not completely heal within a time period of 21 days (osteotomy gap 0.7 mm) in the control group. Thus, an improvement in the healing process can be seen in treated groups. This model only works in female mice since the bone healing process is slower in females than in males (*96*). However, it is not always possible to determine the exact time frames for every phase during the fracture healing process which makes the interpretation much more complex and might impact especially small differences. This might be a reason why the proposed beneficial effect of MIF is not visible indicating a more technical and methodical challenge rather than a biological non-function.

## Materials and Methods

### Single-center retrospective study

In cooperation with the Center for Musculoskeletal Surgery, Charité-Universitätsmedizin Berlin, patient files from patients who were once stationary treated in the hospital during 2012 were screened for ICD-10 classifications M 84.0 (malunion of fracture), M 84.1 (nonunion of fracture) or M 84.2 (delayed union of fracture). Impairment was confirmed by x-rays and patient information as well as patient’s history. Ethical approval for the search algorithm and evaluation sheet was provided by the local ethics committee (EA1/349/13). Due to the retrospective character and anonymization, no consent by the included patients was needed. The selected patient files were additionally reviewed by orthopedic experts before inclusion in the study. Therefore, x-rays and patient information as well as history were re-analyzed in detail. Exclusion criteria were age < 18 years, open fractures and metastases close to the fracture location. Collected data included age, sex, birthday, body height and weight, and fracture related patient’s history including cause, treatment location and complications. In addition, information on lifestyle (e.g. alcoholism, smoking), comorbidities and medications were gathered. The BMI was calculated based on body height and weight. For statistical analysis, the modelling was performed based on univariable and multivariable logistic regression using SPSS V. 22. The first model was built to determine the potential influence of each variable on the fracture healing outcome. The second model served to identify potential confounding factors and verify the independent contribution of variables to the fracture healing outcome.

### Systematic literature review

For the systematic literature review the following search terms were used for a Pubmed based search: (“deferoxamine”[Tiab] OR “DFO”[Tiab] OR “DFX” [Tiab] OR “PHD inhibitor”[Tiab]) AND (“fracture”[Tiab] OR “fracture healing”[Tiab] OR “bone healing”[Tiab] OR “bone regeneration”[Tiab] OR “bone formation”[Tiab] OR “osteotomy”[Tiab]) - Filters activated: Publication date to 2019/02/28. Google scholar was searched in addition with the terms: deferoxamine, bone healing, fracture. The detailed search strategy is comprehensively explained in **figure S2** following the PRISMA guidelines and recommendations from Syrf and Syrcle.

### Study design – In vitro and in vivo studies

The overall hypothesis of the study was that the local application of MIF/DFO in long bone fractures enhances new bone formation (osteoinduction) and can be used to accelerate fracture healing for the treatment and prevention of fracture healing disorders. For the *in vitro* studies, the endpoints were previously defined by hMSC calcification (Alizarin red staining). For the *in vivo* study the primary endpoint was the bone formation rate (bone volume/total volume) as measured via *ex vivo* μCT after 2 weeks. Additional endpoints were defined by histomorphometry. Four time points were determined for additional endpoint measurements. The healing outcome was investigated via *ex vivo* μCT and histology at day 14 and 21. Two operated animals were excluded due to infection in the osteotomy gap and one animal was partially excluded (only included for *ex vivo* μCT) due to an oblique fixation.

*In vitro* studies using hMSCs were performed as proof of concept and possibility to determine an adequate concentration of DFO/MIF to be used *in vivo*. hMSCs were used from 4 to 6 different donors (biological replicates) in at least triplicates per experiment and condition (technical replicates). Data was only excluded if donors failed to differentiate as indicated by a positive control that was carried out on every plate. Only hMSCs that passed characterization were used. Calcification was measured via Alizarin red staining as selected prospectively.

For the *in vivo* study, power analysis was performed prior to animal tests (nQuery) to determine the animal number with the results to use a minimum of 6 animals to attain worthwhile results and was provided in detail with the animal experiment application. All analyses were performed blinded for the experimenter by randomly numbering the animals. Animals were randomized for pairs and treatment groups, although animals in one cage were treated with the same substances.

### hMSC cultivation and calcification assay

Bone marrow was collected from patients undergoing total hip replacement at the Center for Musculoskeletal Surgery, Charité-Universitätsmedizin Berlin. Samples were registered and distributed by the “Tissue Harvesting” Core Facility of the Berlin Institute of Health Center for Regenerative Therapies (BCRT) (**table S3**). Written consents were gathered from all patients. All protocols were approved by the local ethics committee (EA1/012/13) and performed according to the Helsinki Declaration. hMSC isolation, expansion and full characterization (FACS, differentiation) was performed as described previously (*56, 97*). Expansion was done with DMEM plus GlutaMAX (Thermo Fischer Scientific), 10% FCS (PAA Laboratories), 1% Penicillin-Streptomycin (Thermo Fischer Scientific) at 37°C in 5% CO_2_ atmosphere (app. 18% O_2_). Cells were used within passage 4-7. For the calcification assay, hMSCs were transferred to a 96-well plate with a density of 1×10^4^ cells/well, cultivated for 24 h and treated with osteogenic medium (OM) consisting of DMEM, 10% FCS, 1% Penicillin-Streptomycin, 10 mM β-glycerophosphate, 10^−8^ M dexamethasone water-soluble and 0.1 mM L-ascorbic acid-2-phosphate (Sigma Aldrich). Dexamethasone, Deferoxamine mesylate salt (DFO; Sigma Aldrich) and MIF (lab own production) were supplemented in different concentrations to the medium. Medium was changed weekly. For the release studies OsteoDiff (Miltenyi Biotech) was used supplemented with 1% Penicillin-Streptomycin. For Alizarin red staining cells were fixed with 4% formaldehyde (15 min RT; Carl Roth), washed twice with PBS and stained with 0.5 % Alizarin Red (Sigma Aldrich) in H_2_O_dest_ (pH 4) for 10 min at RT followed by 4 washing steps with H_2_O_dest_ and application of 10% cetylpyridinium chloride solution (AppliChem) for 30 min at RT. Supernatants were transferred to a new 96 well plate and measured with a Synergy HT plate reader (BioTek Instruments) at a wavelength of 562 nm (reference wavelength 630 nm) for quantification.

### Animals, housing and osteotomy

Female C57BL/6N mice (12 weeks; body weight 20 – 25 g; Charles River Laboratories) were housed in the Charité animal facility (FEM; semi-sterile - outside the SPF barrier) in pairs in Euro standard Type II clear-transparent plastic cages and kept under obligatory hygiene standards monitored according to the FELASA standards. Nesting material was provided in sufficient amount while pipes and houses were withdrawn after osteotomy to avoid possible entanglement with the used external fixator. Food and water were available ad libitum and the temperature was (20 ± 2 °C) controlled with a 12 h light/dark period and a humidity of 45-50%.

All experiments were carried out with ethical permission according to the policies and principles established by the Animal Welfare Act, the National Institutes of Health Guide for Care and Use of Laboratory Animals, and the National Animal Welfare Guidelines, the ARRIVE guidelines and were approved by the local legal representative animal rights protection authorities (Landesamt für Gesundheit und Soziales Berlin: G 0111/13, 0039/16). Pain management and osteotomy were performed as described in detail previously (*56, 57, 98, 99*). In short, for analgesia a buprenorphine injection (0.1 mg/kg; Temgesic, RB Pharmaceuticals Limited) *s.c.* was given prior to the surgery and Tramadol (0.1 mg/ml; Drops, Grünenthal GmbH) was applied with the drinking water for the first 3 post-operative days. Anesthesia was conducted with isoflurane and O_2_ supplementation and mice were prepared with eye ointment (Bayer Pharma AG) and clindamycin *s.c.* (0.02 ml; Ratiopharm GmbH). Osteotomy with an external fixator (MouseExFix, RISystem) was performed at the left femur creating a 0.70 mm osteotomy gap with a Gigli wire saw. The osteotomy gap was filled with PBS-soaked ACS (control; Lyostypt, B. Braun) or MIF and/or DFO solved in PBS applied on the ACS (treatment groups; 100 ng/ml and/or 250 μM, respectively) (*56, 98*). After skin closure, mice received warmed NaCl (0.2 ml) *s.c* returned to their cages with a prepared nest and an infrared radiator. Animals were euthanized via cervical dislocation after 3, 7, 14 and 21 days in deep anesthesia (no deep pain perception) after intracardial blood collection. Osteotomized femora were collected and either fixed with 4% paraformaldehyde (PFA; Electron Microscopy Sciences) for 6-8 h at 4°C.

### Ex vivo micro computed tomography (μCT)

PFA-fixed femora were treated with an ascending sucrose solution (10%, 20%, 30%) for 24 h, respectively at 4°C. Scanning of 191 slices was performed after removal of the pins and external fixator with an isotropic voxel size of 10.5 μm (70 KVp, 114 μA; SCANCO μCT Viva 40), aligned scan axis along the diaphyseal axis of the femora and 3D reconstruction and analyses were performed using the provided software package as described previously and applying a fixed global threshold of 240 mg HA/cm^3^ for the automatic 3D callus tissue analysis (*56, 98*). Nomenclature and analysis were conducted in accordance with published recommendations (*100*).

### Histology and immunofluorescence

After μCT scanning, femora were cryo-embedded without decalcification according to the Kawamoto *et al*. method (*101*). For Movat’s pentachrome staining slices (7 μm) were air dried for 30 min, fixed with 4 % PFA for 10 min and washed with H_2_O_dest_ for 5 min. The staining procedure was conducted using a protocol already been published (*56, 102*). The staining results allowed to distinguish between different tissues: mineralized bone or mineralized cartilage appear yellow, hyaline cartilage green, cytoplasm reddish, cell nuclei blue-black and the surrounding muscles are colored in reddish. When combined with Von Kossa staining – the following staining steps were conducted before the Movat’s pentachrome staining: 3% (w/v) silver nitrate solution (10 min), washing step with H_2_O_dest_, sodium carbonate formaldehyde solution (2 min), washing step with tap water, 5% (w/v) sodium thiosulphate solution (5 min), washing step with tap water and H_2_O_dest_. Images were taken with a light microscope (Leica) in a 2.5x magnification and the program Axiovision (Carl Zeiss Microscopy). The Acid Phosphatase, Leukocyte (TRAP) Kit (Sigma Aldrich) was used to stain for TRAP. Manufacturer’s instructions were followed, and pictures were taken with a light microscope (Leica).

For immunofluorescence staining the following primary antibodies: CD31/PECAM-1 (goat polyclonal unconjugated, AF2628, R&D Systems, 1:100), Emcn (V.7C7 unconjugated, sc-65495, 1:100), F4/80 (Cl:A3-1 unconjugated, MCA497G, 1:400), Osx (rabbit polyclonal, sc-22536-R, 1:200) and secondary antibodies (all Thermo Fischer Scientific; 1:500): anti-rat conjugated AF594 (A21209), anti-rabbit conjugated AF488 (A21206), anti-rat conjugated AF647 (A21247), anti-goat conjugated AF647 (A21447) or anti-goat A568 (A11057) were used. Staining procedure was performed in a wet section as published earlier (*56, 57*). Pictures were taken with a fluorescence microscope BZ 9000 (Keyence). Image analysis was performed with ImageJ (*56, 57, 98*).

### DFO release assay

In order to measure the DFO concentration in the supernatant we established the method described by Fielding and Brunström 1964 (*103*). The method is based on the ability of DFO to bind iron and therefore reduces ferric chloride (Fe^3+^) to ferrioxamine (red-brown compound). Measurements were taken including all components (citric acid and Na_2_HPO_4_) and a ferric chloride concentration of 1.5 mg/100 ml at a wavelength of 450 nm (Synergy HT plate reader, BioTek Instruments). Release kinetic experiments were performed in PBS. Hydrogel formation was performed as described in detail before (*104*). 100 μl of 20% dPG or dPGS were mixed with 10.000 μM DFO and polymerized for 1h at 37°C. For release kinetic and transfer assay 200 μl PBS were added and supernatants were collected every 24h. For transfer assay to HEK 293 cells supernatant was collected after 24h and mixed with normal expansion medium (1:10). HEK 293 were treated for 24h before collecting protein and performing HIF-1α western blot as described before (*105*). For co-cultivation with hMSCs, hydrogels were polymerized in 24-well transwell inserts (Sarstedt; polyethylene terephthalate membrane with 8 μm pore size). After polymerization, hydrogels were treated for 12h with osteogenic medium at 37°C for equilibration and transferred to a 24-well plate seeded with hMSCs. Co-cultivation was performed for 14 days with supplementation of osteogenic medium. Alizarin red staining was used to visualize calcification of hMSCs.

### Statistical analysis

Statistical analysis was carried out with GraphPad Prism V.8 software. All values from *in vitro* assays were expressed as the mean ± SD or SEM when measured in > duplicates and all values from animal experiments are depicted as median ± ranges (box and whiskers plot with individual data points). Kruskal Wallis test with Dunn’s multiple comparison test and Wilcoxon-signed rank test were applied in case of lack of Gaussian distribution that was tested before via Kolmogorov-Smirnov test. A *p-value* <*0.05* was considered as statistically significant. In some cases, statistical trends are indicated with a hashtag (#) when biologically relevant.

## Acknowledgments

The authors would like to thank Gabriela May, Manuela Jakstadt, Gabriela Korus, Marie-Christin Weber and Mario Thiele for excellent technical assistance. Additionally, we like to acknowledge Peter Schlattmann (University of Jena, Germany) for supporting logistic regression analysis. Moreover, Anastasia Rakow and Paul Simon served as orthopedic expert team to re-analyze the cases included in the retrospective study. FACS analyses were performed together with the Core Facility at the German Rheumatism Research Centre. Bone-marrow was provided from the “Tissue Harvesting” Core Facility of the BCRT. This study is a part of Annemarie Lang’s PhD thesis as published under https://refubium.fu-berlin.de/handle/fub188/26365. The retrospective study was performed within the doctoral dissertation of Sara Fügener (now Helfmeier) - https://refubium.fu-berlin.de/handle/fub188/3611?show=ull.

## Author contributions

K.S.B., G.N.D., M.L., T.G. and F.B. conceived and supervised the research. S.H., A.L., P.H., C.P. and F.B. designed the retrospective study, performed data collection and quality control and performed analysis. A.L., K.S.B. an T.G. designed and performed in vitro and animal experiments. A.L., J.S., A.K., V.S., A.W. A.D. and M.P. performed analysis. A.L., T.G., J.R., S.H.-S., R.H. planned, supervised or performed release assays. A.L., F.B., T.G. and K.S.B. wrote the paper. A.E.H., G.N.D. and M. L. revised manuscript.

## Data and materials availability

All data associated with this study are present in the paper or the Supplementary Materials.

## Supplementary Materials

## 1. Retrospective study - Detailed information and additional data

**Figure S6:**
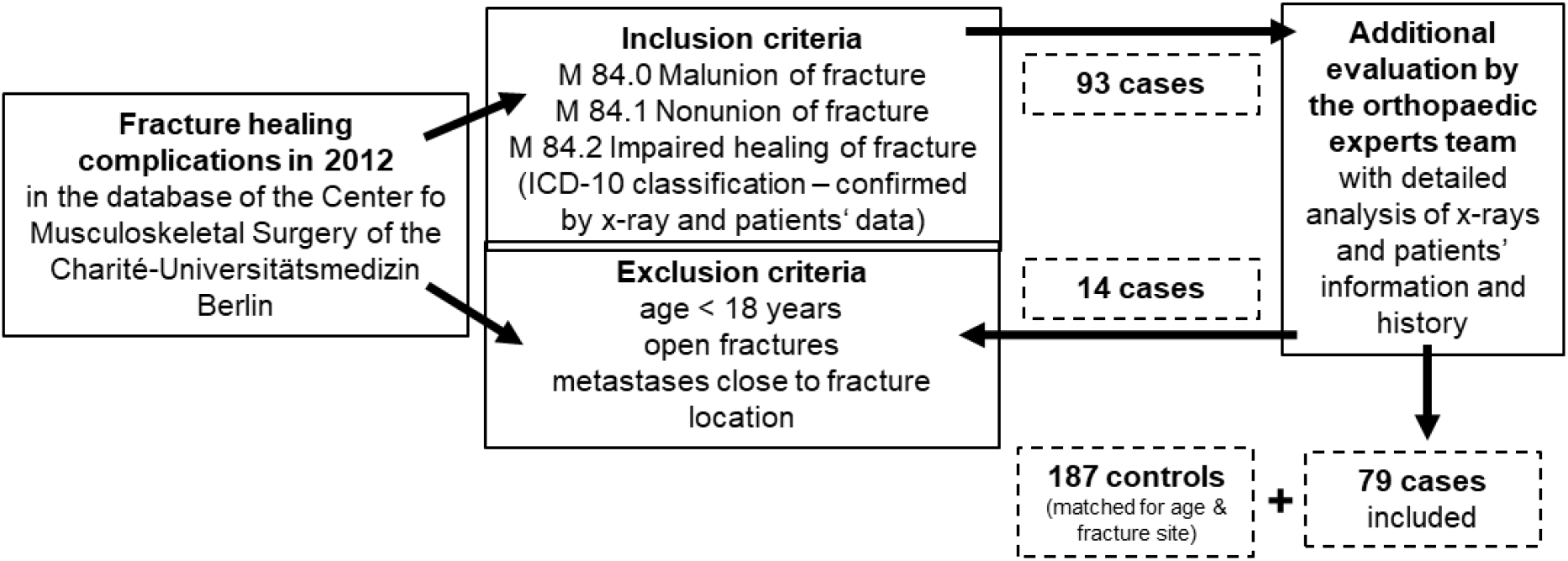
Search strategy of the retrospective study including inclusion and exclusion criteria.

**Table S1:** Detailed characterization of the 79 included cases.

|  | Total number | in % |
| --- | --- | --- |
| **ICD-10 classification** |  |  |
| M 84.0 | 12 | 15.2 |
| M 84.1 | 61 | 77.2 |
| M 84.2 | 6 | 7.6 |
| **Fracture localisation** |  |  |
| Humerus | 15 | 19.0 |
| Radius, ulna | 12 | 15.2 |
| Femur | 20 | 25.3 |
| Tibia, fibula | 11 | 13.9 |
| Clavicle | 7 | 8.9 |
| Scaphoid | 5 | 6.3 |
| Vertebrae | 4 | 5.1 |
| Os sacrum | 1 | 1.3 |
| Patella | 1 | 1.3 |
| Metatarsal | 3 | 3.8 |

**Table S2:** Descriptive analysis.

|  | Controls in % | Cases in % |
| --- | --- | --- |
| **Age** |  |  |
| 18 - 39 | 26.2 | 30.4 |
| 40 - 59 | 36.9 | 34.2 |
| > 60 | 36.9 | 35.4 |
| **Gender** |  |  |
| male | 44.9 | 53.2 |
| female | 55.1 | 46.8 |
| **BMI** |  |  |
| < 20 | 9.9 | 6.5 |
| 20 - 25 | 40.7 | 41.6 |
| > 25 | 49.4 | 51.9 |
| **Alcohol abuse** |  |  |
| no | 85.6 | 97.3 |
| yes | 14.4 | 2.7 |
| **Smoking** |  |  |
| no | 71.9 | 62.2 |
| yes | 28.1 | 37.8 |
| **Glucocorticoids** |  |  |
| no | 97.3 | 93.7 |
| continuously | 1.6 | 6.3 |
| temporarily | 1.1 | 0 |
| **NSAIDs** |  |  |
| no | 84.5 | 86.1 |
| continuously | 2.7 | 7.6 |
| ASS-100 mg | 12.8 | 6.3 |
| **Rheumatoid Arthritis** |  |  |
| no | 99.5 | 93.7 |
| yes | 0.5 | 6.3 |
| **Osteoporosis** |  |  |
| no | 90.8 | 89.9 |
| yes | 9.5 | 10.1 |
| **Arterial Hypertension** |  |  |
| no | 66.5 | 57.0 |
| yes | 33.5 | 43.0 |
| **Diabetes Type 2** |  |  |
| no | 88.1 | 88.6 |
| yes | 11.9 | 11.4 |

## 2. Systematic literature review – Search strategy

**Figure S7:**
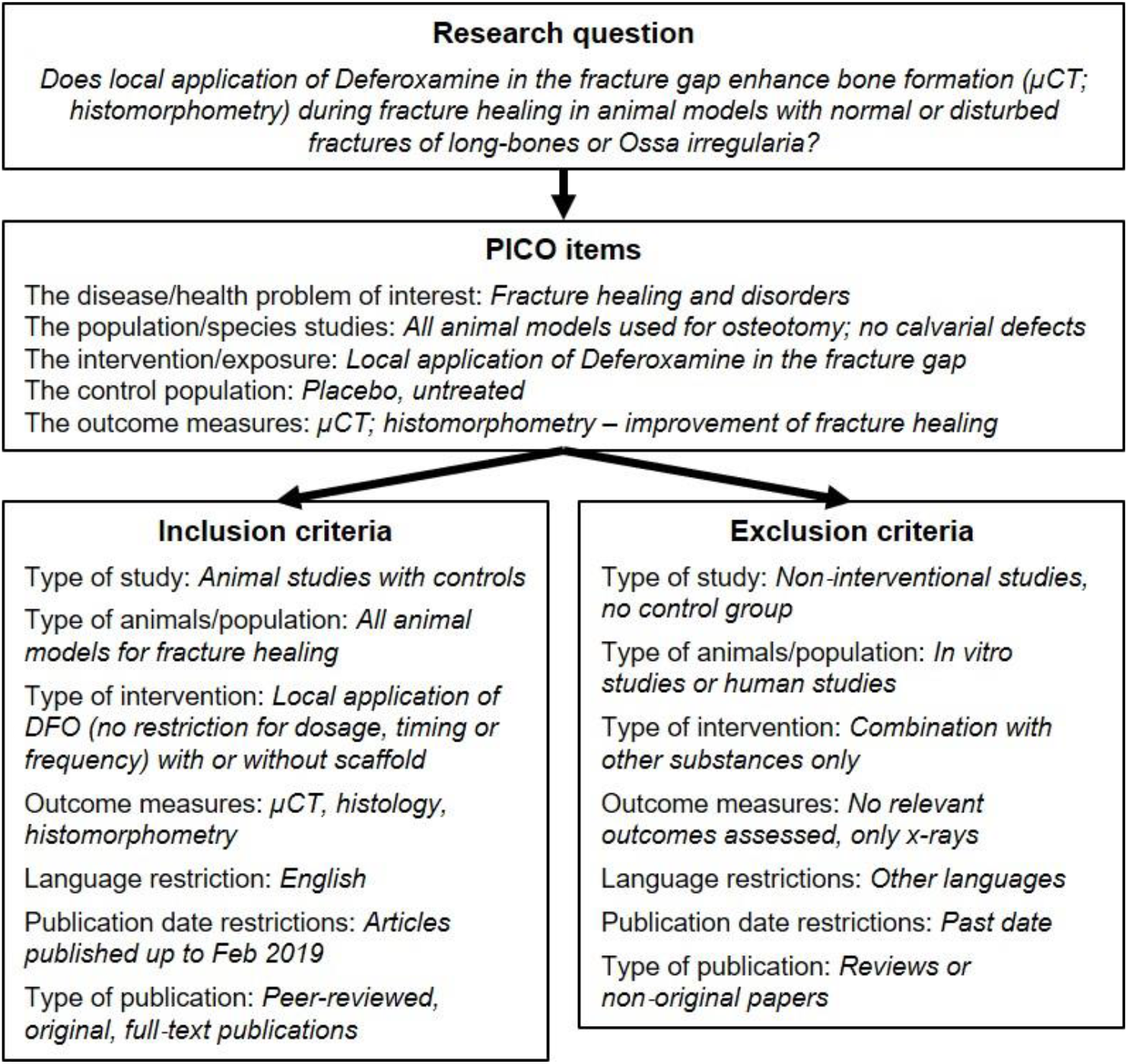
Search strategy of the systematic literature review in accordance with the PRISMA guidelines and recommendations from Syrf and Syrcle.

**Figure S8:**
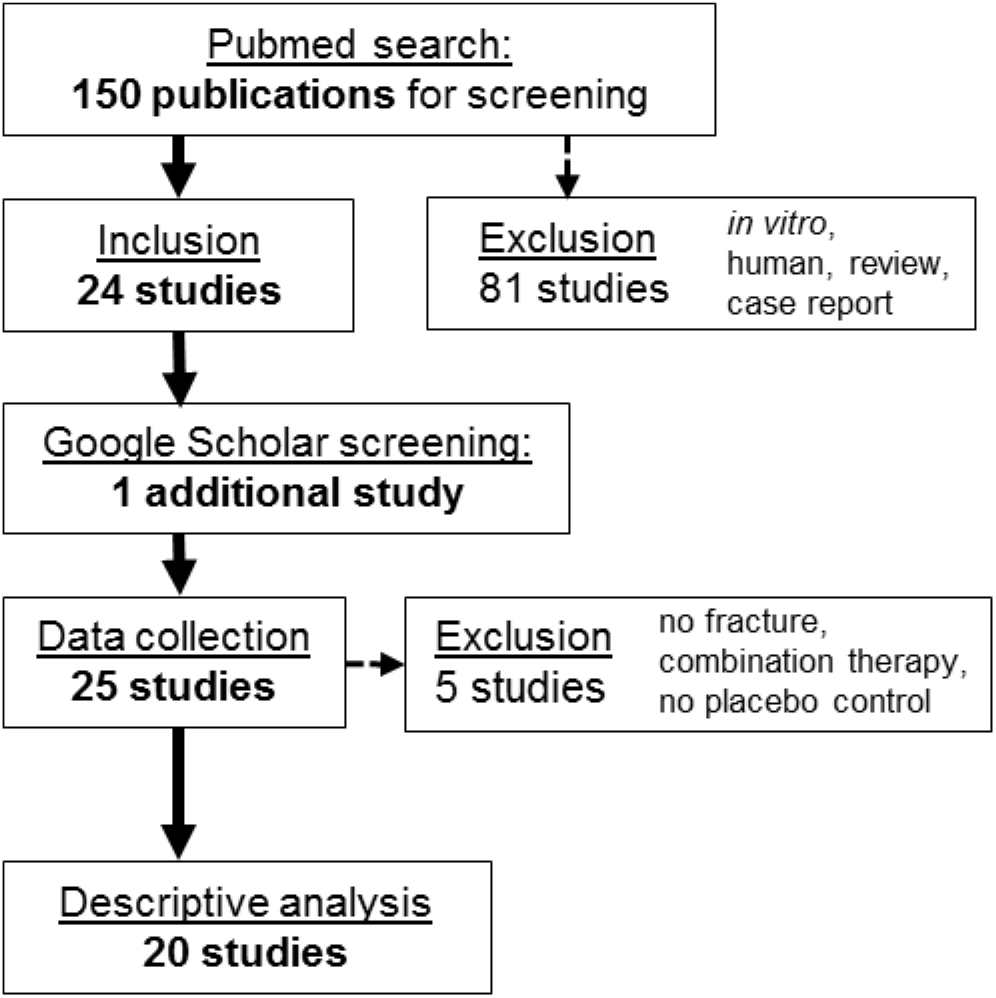
Flow diagram of the systematic literature review resulting in the inclusion of 20 studies.

## 3. *In vitro* and *in vivo* studies – Additional data

**Figure S9:**
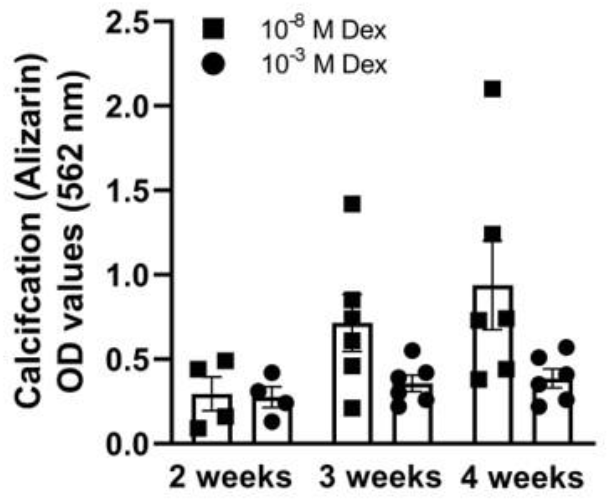
In vitro model pre-testing. Determination of the conditions and testing period of the calcification assay for further titration experiments under normoxic conditions (N = 4-6; > triplicates). Bar graphs show mean ± SEM and individual data points.

**Figure S10:**
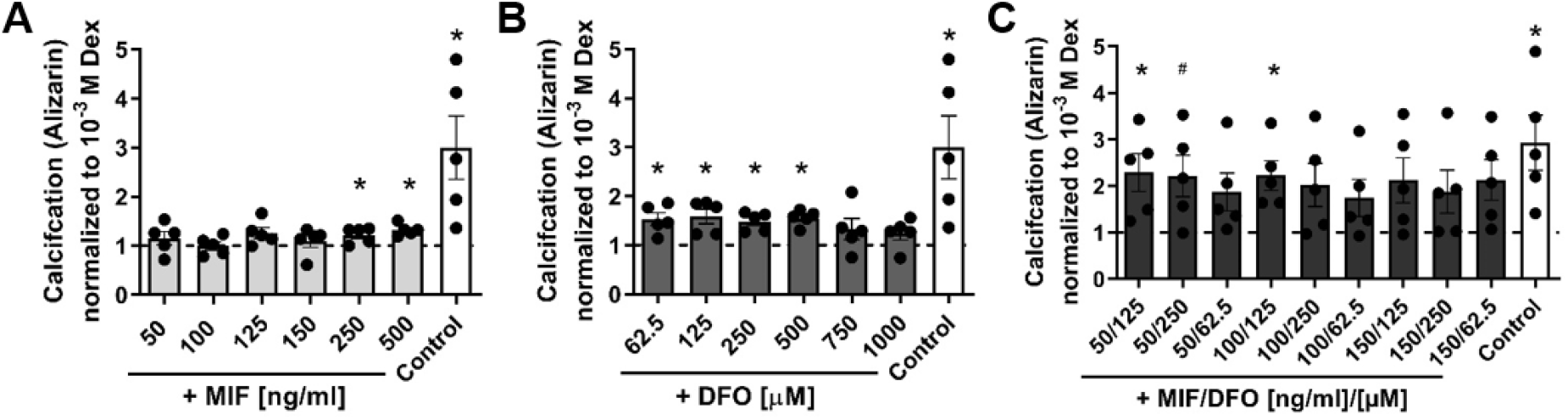
Titration of different MIF and DFO concentrations alone (A, B) and in combination (C). hMSCs were cultivated under normoxia for 4 weeks with addition of osteogenic medium (OM). OD values gained after Alizarin red staining were normalized to Dex 10^−3^ M (N = 4-6; > triplicates). Bar graphs show mean ± SEM and individual data points. One sample t-test was used to determine statistical significance towards the hypothetical value of 1. p-values are indicated with ^#^P < 0.07; *P < 0.05.

**Figure S11:**
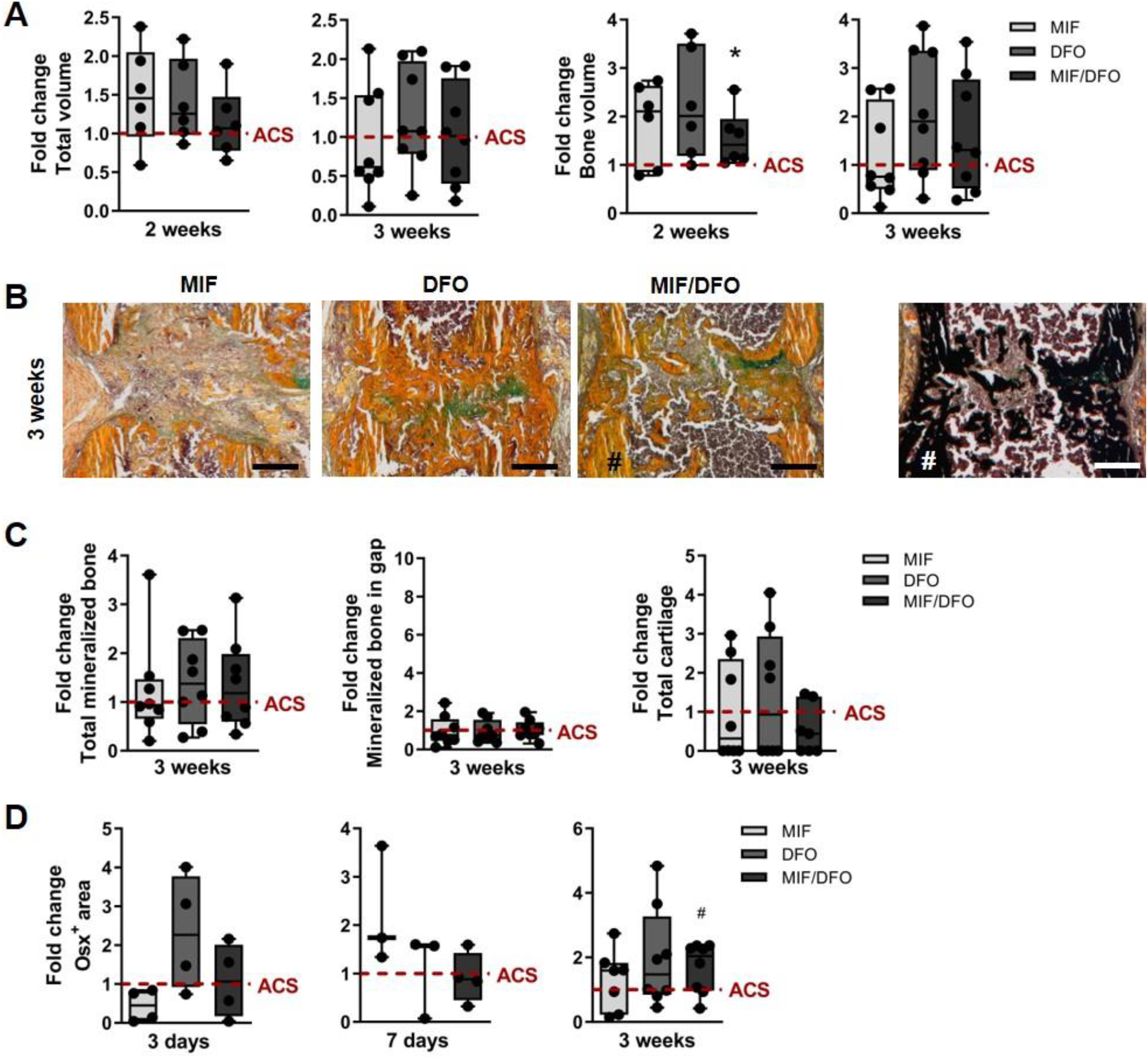
Bone regeneration in a delayed healing model after single dose of MIF or/and DFO – additional data. (A) MicroCT quantification of total volume and bone volume at 2 weeks and at 3 weeks post-osteotomy normalized to the median of the ACS group (indicated as dotted line = 1). (B) Representative images of Movat’s pentachrome staining for each treatment group. yellow-mineralized bone/scaffold; green – cartilage; magenta – bone marrow. Representative images for ^#^MIF/DFO, 3 weeks of von Kossa combined with Movat’s pentachrome staining to show the distinction between mineralized bone and residual scaffold. (C) Histomorphometry of Movat’s pentachrome staining using ImageJ. Data were normalized to the median of the ACS group (indicated as dotted line = 1). (D) Quantification of Osx staining at 3, 7 days and 3 weeks. Data are shown as box plots with the median as horizontal line, interquartile range as boxes, minimum/maximum as whiskers and individual data points. Wilcoxon signed rank test was applied to determine difference against the ACS control group (hypothetical value = 1) and Kruskal Wallis test with Dunn’s multiple comparison test was used to compare groups. ^#^P < 0.07; *P < 0.05.

**Figure S12:**
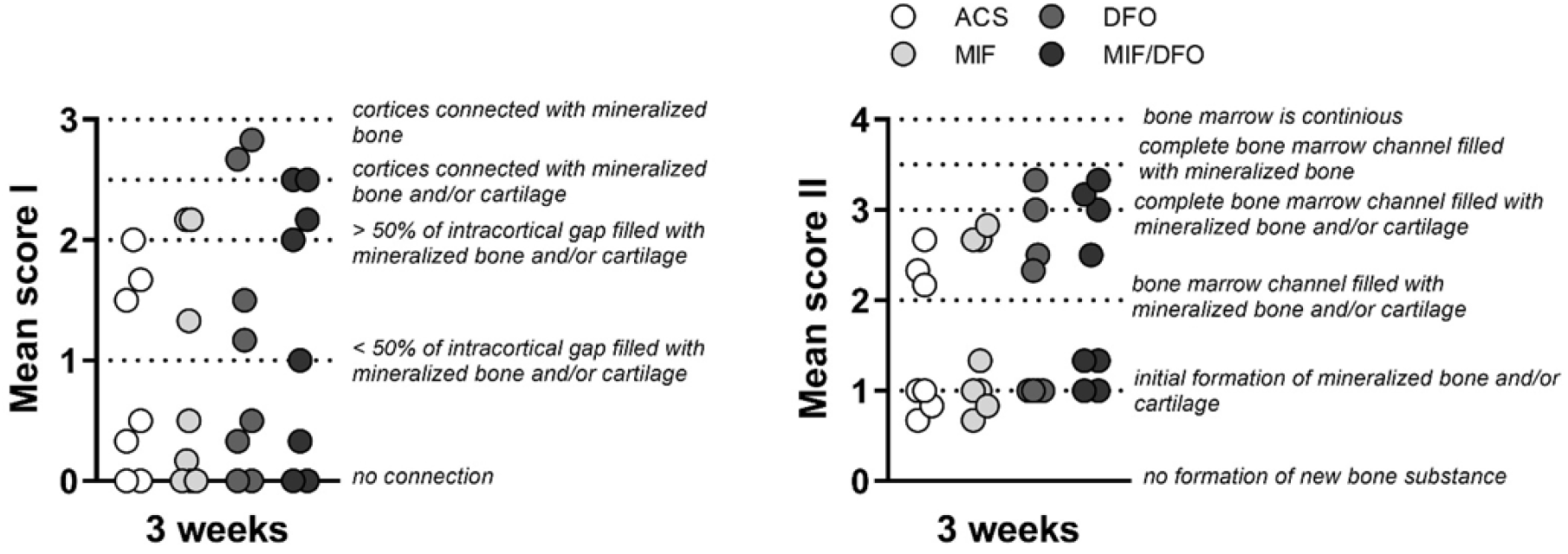
Results of two qualitative scores that were performed with all slides (Movat’s pentachrome staining) by 3 independent experimenters. Score criteria are specified in the graphs. Mean score I focused in the intracortical area indicating a bridging between the cortices. Mean score II focus in the filling of the fracture gap/ bone marrow area between cortices. The qualitative score was aimed to underline subjective findings and the quantification via histomorphometry.

**Figure S13:**
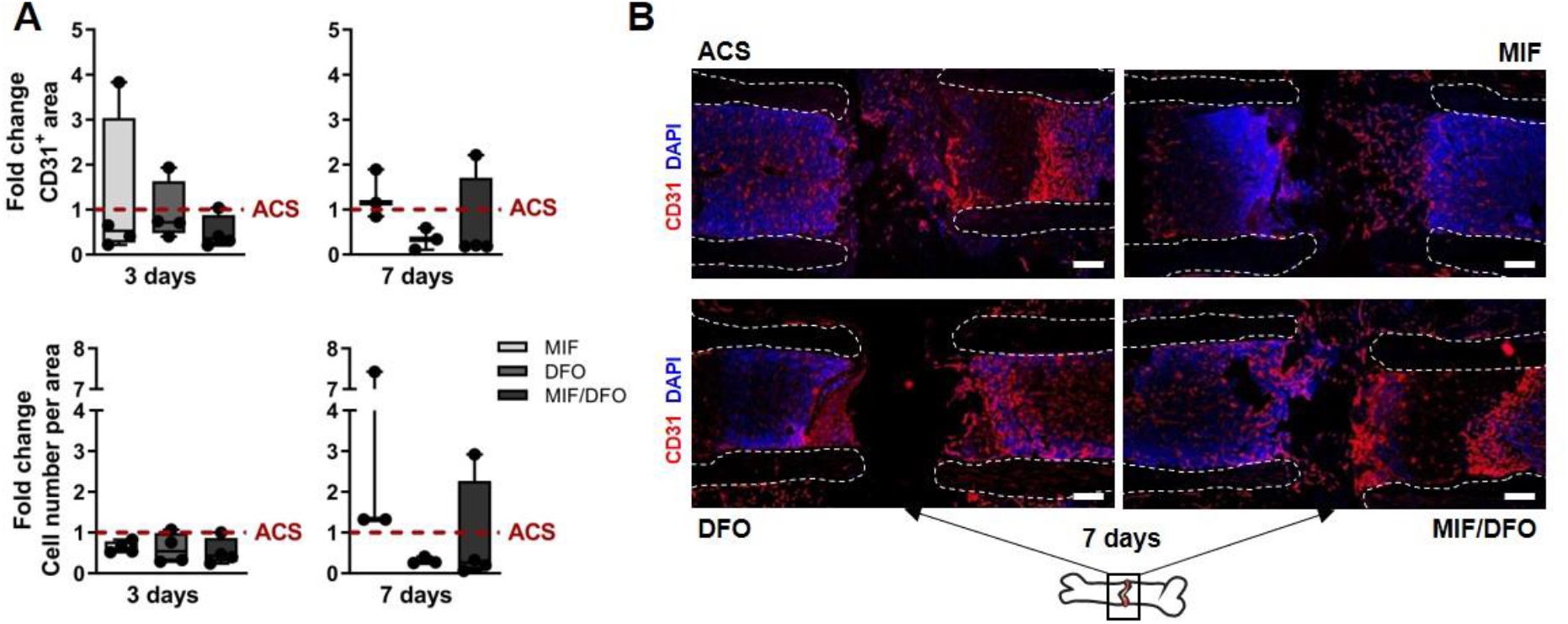
Revascularization in a delayed healing model under MIF, DFO and MIF/DFO treatment – additional data. (A) Quantification CD31^+^ stained areas and cell numbers per area normalized to the median of the ACS group (indicated as dotted line = 1) and (B) corresponding representative images for day 3 and 7 (N = 3-4). White dotted lines indicate cortices. Schematic bone indicates alignment of images. Scale bars = 200μm. Data are shown as box plots with the median as horizontal line, interquartile range as boxes, minimum/maximum as whiskers and individual data points. Wilcoxon signed rank test was applied to determine difference against the ACS control group (hypothetical value = 1) and Kruskal Wallis test with Dunn’s multiple comparison test was used to compare groups.

**Figure S14:**
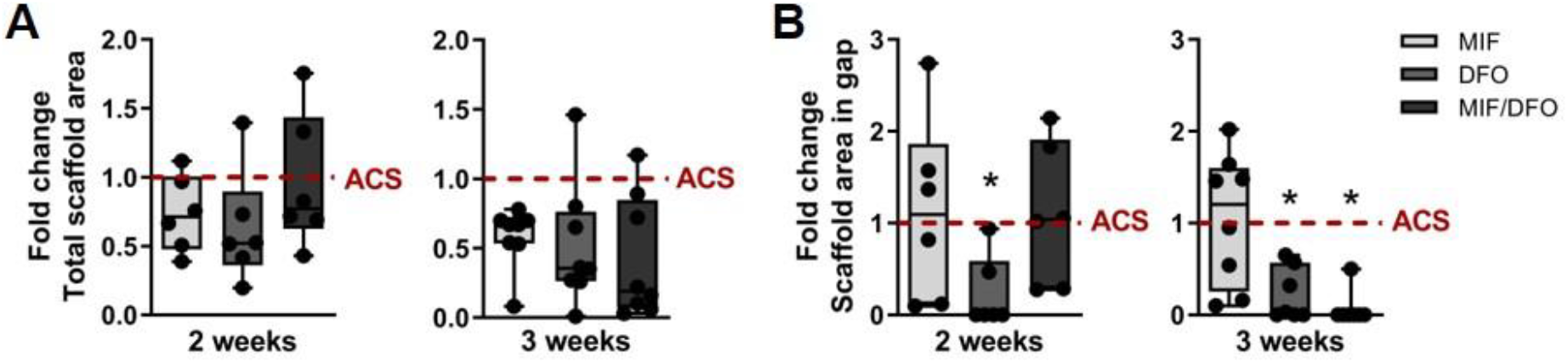
Presence of F4/80+ macrophages and TRAP+ cells within the fracture gap – additional data. (A) Quantified total scaffold areas and (B) in the gap after 2 and 3 weeks normalized to the median of the ACS group (indicated as dotted line = 1). (N = 6-8). Data are shown as box plots with the median as horizontal line, interquartile range as boxes, minimum/maximum as whiskers and individual data points. Wilcoxon signed rank test was applied to determine difference against the ACS control group (hypothetical value = 1) and Kruskal Wallis test with Dunn’s multiple comparison test was used to compare groups. *P < 0.05.

**Table S3:**
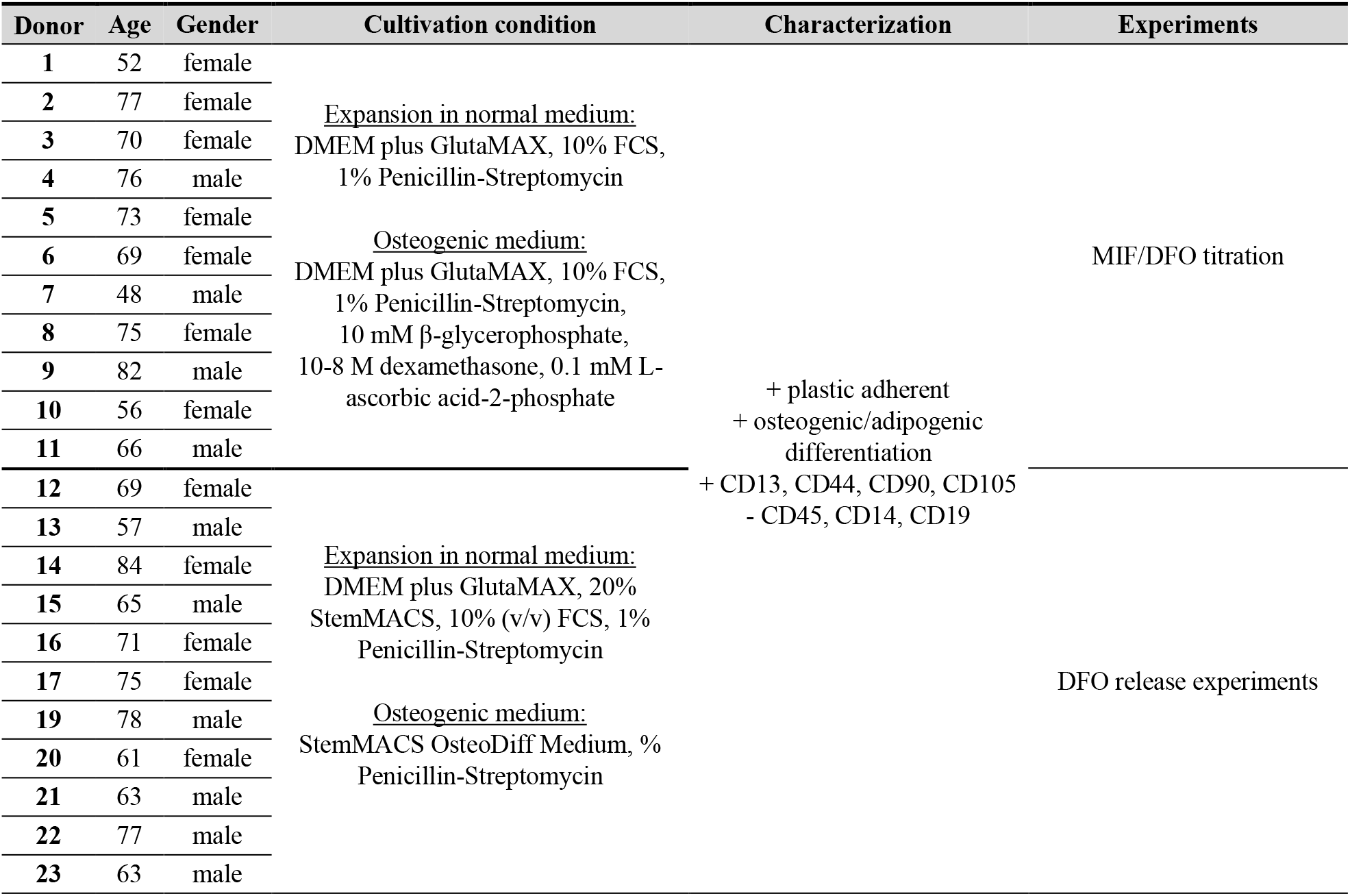
List of hMSCs that were used in the study.

